# Oscillatory traveling waves reveal predictive coding abnormalities in schizophrenia

**DOI:** 10.1101/2023.10.09.561490

**Authors:** Andrea Alamia, Dario Gordillo, Eka Chkonia, Maya Roinishvili, Celine Cappe, Michael H. Herzog

## Abstract

The computational mechanisms underlying psychiatric disorders are hotly debated. One hypothesis, grounded in the Bayesian predictive coding framework, proposes that schizophrenia patients have abnormalities in encoding prior beliefs about the environment, resulting in abnormal sensory inference, which can explain core aspects of the psychopathology, such as some of its symptoms. Here, we tested this hypothesis by identifying oscillatory traveling waves as neural signatures of predictive coding. By analyzing a EEG dataset. comprising 146 schizophrenia patients and 96 age-matched healthy controls, we found that schizophrenia patients have stronger top-down alpha-band traveling waves compared to healthy controls during resting state, reflecting stronger precise priors at higher levels of the predictive processing hierarchy. Conversely, we found stronger bottom-up alpha-band waves in schizophrenia patients during a visual task reflecting an alteration of lower sensory priors. Our results yield a novel spatial-based characterization of oscillatory dynamics in schizophrenia, considering brain rhythms as traveling waves and providing a unique framework to study the different components involved in a predictive coding scheme. Altogether, our findings significantly advance our understanding of the mechanisms involved in fundamental pathophysiological aspects of schizophrenia, promoting a more comprehensive and hypothesis-driven approach to psychiatric disorders.

**Significance:** We provide novel evidence favoring the Bayesian predictive coding interpretation of schizophrenia. Relying on computational and experimental works that characterized electrophysiological correlates of predictive processes, we investigate oscillatory traveling waves in EEG data of 146 schizophrenia patients and 96 age-matched healthy controls. Our results reveal stronger top-down alpha-band traveling waves in schizophrenia patients, reflecting an increase in the precision of high-level priors. On the other hand, we observed an increase in bottom-up alpha-band waves during a visual task, in line with the proposed reduction in precision of low-level sensory priors. Our findings suggest that traveling waves’ analysis is a versatile technique to probe predictive processing mechanisms in different cognitive processes. Impairments in this mechanism may underlie perceptual alterations as well as the pronounced clinical symptoms of schizophrenia. The strong hypothesis-driven nature of our results accentuates the relevance of our findings.

## Introduction

Schizophrenia is a severe mental disorder that affects about one percent of the world’s population (McCutcheon et al., 2020). Schizophrenia is characterized by a large range of psychotic symptoms as well as by strong impairments in mental functioning, including perception, cognition, and personality.

Numerous hypotheses and mechanisms have been proposed to explain these abnormalities. One hypothesis is that schizophrenia patients dysfunctionally update their cognitive world model, usually described within the framework of Bayesian inference and predictive coding (Corlett et al., 2009; Krystal et al., 2017). According to this framework, perception combines incoming sensory evidence with prior information, i.e., beliefs about the world (figure 1A). However, experimental evidence from behavioral studies is mixed, including studies showing that patients rely more on prior beliefs than on sensory information (Cassidy et al., 2018; Powers et al., 2017), studies which found stronger reliance on sensory information in the patients (Stuke et al., 2019; Weilnhammer et al., 2020), and even studies which found intact processing (Choung et al., 2022; Kaliuzhna et al., 2019; Tibber et al., 2013).

**Figure 1.**
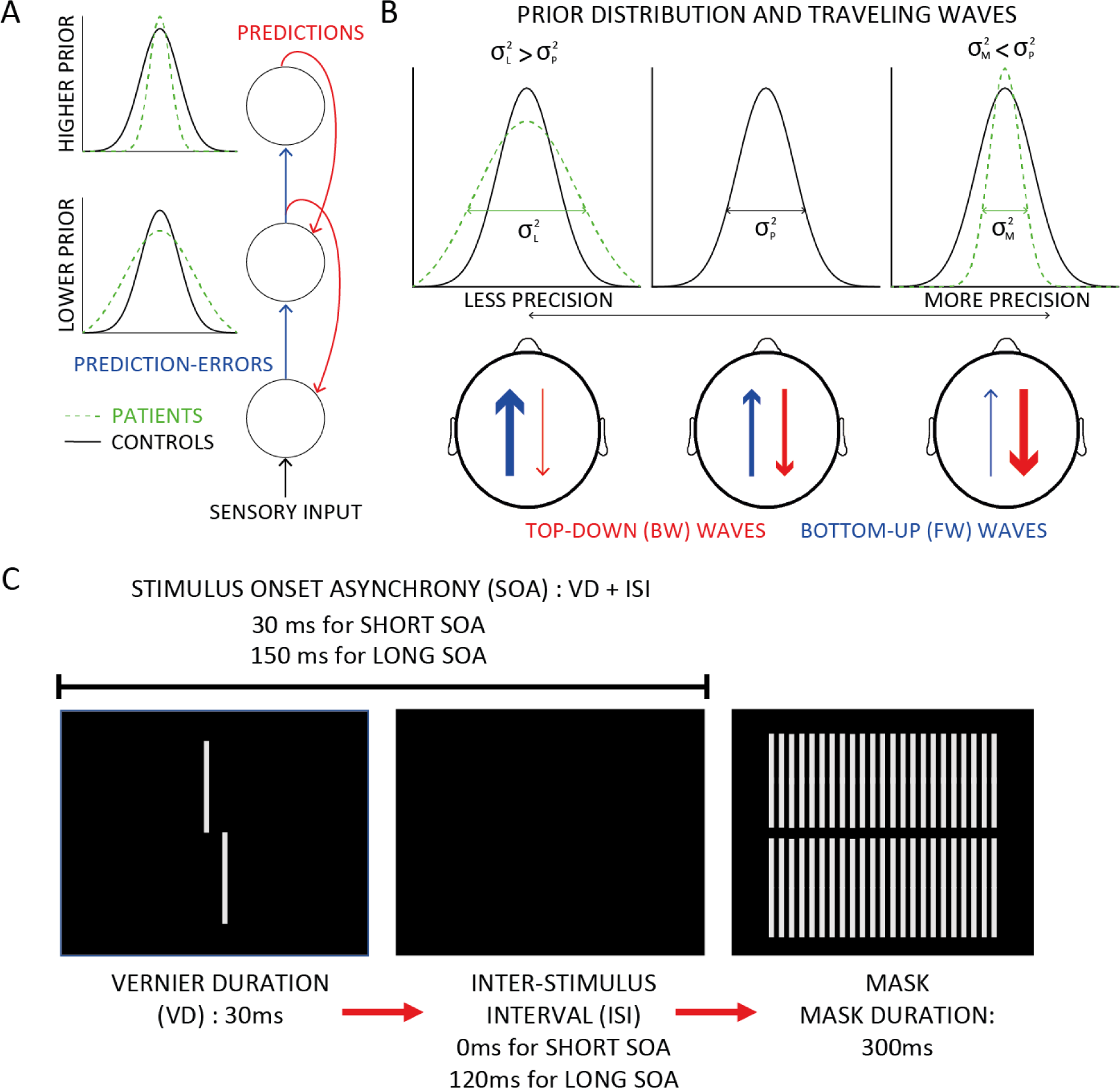
Predictive coding and traveling waves. A) In the Bayesian predictive coding perspective, predictions are generated by prior distributions in higher brain regions, prediction-errors are computed to update the prior based on the sensory evidence (i.e., the likelihood). Recently, it was proposed that hierarchical-specific abnormalities of the priors’ precision in schizophrenia, have better precision in higher areas and worse precision in lower, sensory-related areas (Corlett et al., 2019). B) Considering backward (BW) and forward (FW) waves as proxies of predictions and prediction-errors, respectively (Alamia & VanRullen, 2019), one would expect different patterns of traveling waves depending on the precision of the prior: more precise prior (rightmost panel) generate stronger predictions, and in turn stronger backward waves, whereas less precise prior information (leftmost pattern) generates inaccurate predictions, hence higher prediction-errors, reflected by stronger forward waves. C) In our experiment, the target consisted of a Vernier, i.e., two vertical lines slightly offset horizontally either to the left or right (as shown). Participants were instructed to indicate the offset direction. A grating mask was presented afterwards, making the discriminability of the offset spatially and temporally challenging.

Basic sensory impairments and hallucinations may however rely on abnormalitites at different levels of predictive processing. Hence, it has been proposed that schizophrenia patients weigh the prior information more strongly in hierarchically higher-regions, but rely more on sensory information in lower hierarchical regions (Corlett et al., 2019; Sterzer et al., 2018). These hierarchical-specific alterations in the priors would explain both impairments in basic sensory processing and also more complex phenomena such as hallucinations or delusions (Friston et al., 2016; Krystal et al., 2017), and may therefore explain the mixed experimental results.

Here, we tested the hypothesis that there are hierarchical-specific alterations in predictive processing from a *neurophysiological* perspective. In recent work, it was shown that oscillatory alpha-band (8-12Hz) traveling waves are neural signatures of predictive coding (Alamia & VanRullen, 2019; Arnal & Giraud, 2012; Bastos et al., 2012, 2015; Friston, 2019; Michalareas et al., 2016). In these studies (Alamia & VanRullen, 2019; Pang et al., 2020), neural activity was measured along the central midline of electrodes (Oz-Fz) to determine how oscillations propagate as traveling waves from occipital to frontal areas (forward waves) or in the opposite direction (backward waves). Based on a model, implementing predictive coding under minimal assumptions (Alamia & VanRullen, 2019), the authors proposed that forward traveling waves encode sensory information and prediction-errors (i.e., the difference between top-down predictions and the actual activity), while backward waves carry the prior information (Rao & Ballard, 1999).

A pharmacological study provided additional evidence for the relationship between traveling waves and predictive coding using a serotonergic psychedelics drug (i.e., N, N-Dimethyltryptamine, DMT; Alamia et al., 2020). According to a recently proposed hypothesis (Carhart-Harris & Friston, 2019), psychedelics act on the high-level prior distributions in the brain, decreasing their precision (defined as the inverse of the variance, figure 1B). As a consequence, there is an increase in forward waves carrying prediction-errors (figure 1B, left panel). Indeed, after the intake of DMT, the authors observed a decrease and an increase in the alpha-band traveling waves propagating top-down and bottom-up, respectively. Here, we used the very same analysis to test whether patients rely more on prior beliefs than on sensory evidence during eyes-closed, resting-state EEG and a visual backward masking task. Following the Bayesian framework, we expect stronger alpha band backward waves during resting state in schizophrenia patients than in healthy controls, due to more precise high-order priors; on the other hand, we expect stronger forward alpha-band waves during a visual task, reflecting an increase in the weighting of the sensory information. (figure 1B). We analyzed a large EEG dataset comprising 146 schizophrenia patients and 96 age-matched healthy controls.

## Results

We re-analyzed EEG data from previous studies with resting state data (eyes closed; da Cruz et al., 2020; Gordillo et al., 2023) and a visual backward masking task (da Cruz, Shaqiri, et al., 2020; Garobbio et al., 2021). We quantified brain oscillations along the central electrodes’ mid-line (from Oz to Fz, figure 2), as in our previous work (Alamia & VanRullen, 2019; Pang (庞兆阳) et al., 2020). We considered sliding time windows of 1 and 0.5 seconds from two different datasets, one eyes-closed resting state and one visual backward masking (VBM) task (see Methods and figure 2). First, for all time windows, we create 2D maps by stacking the signals from the seven electrodes, obtaining images with time and electrodes as axes (figure 2).

**Figure 2.**
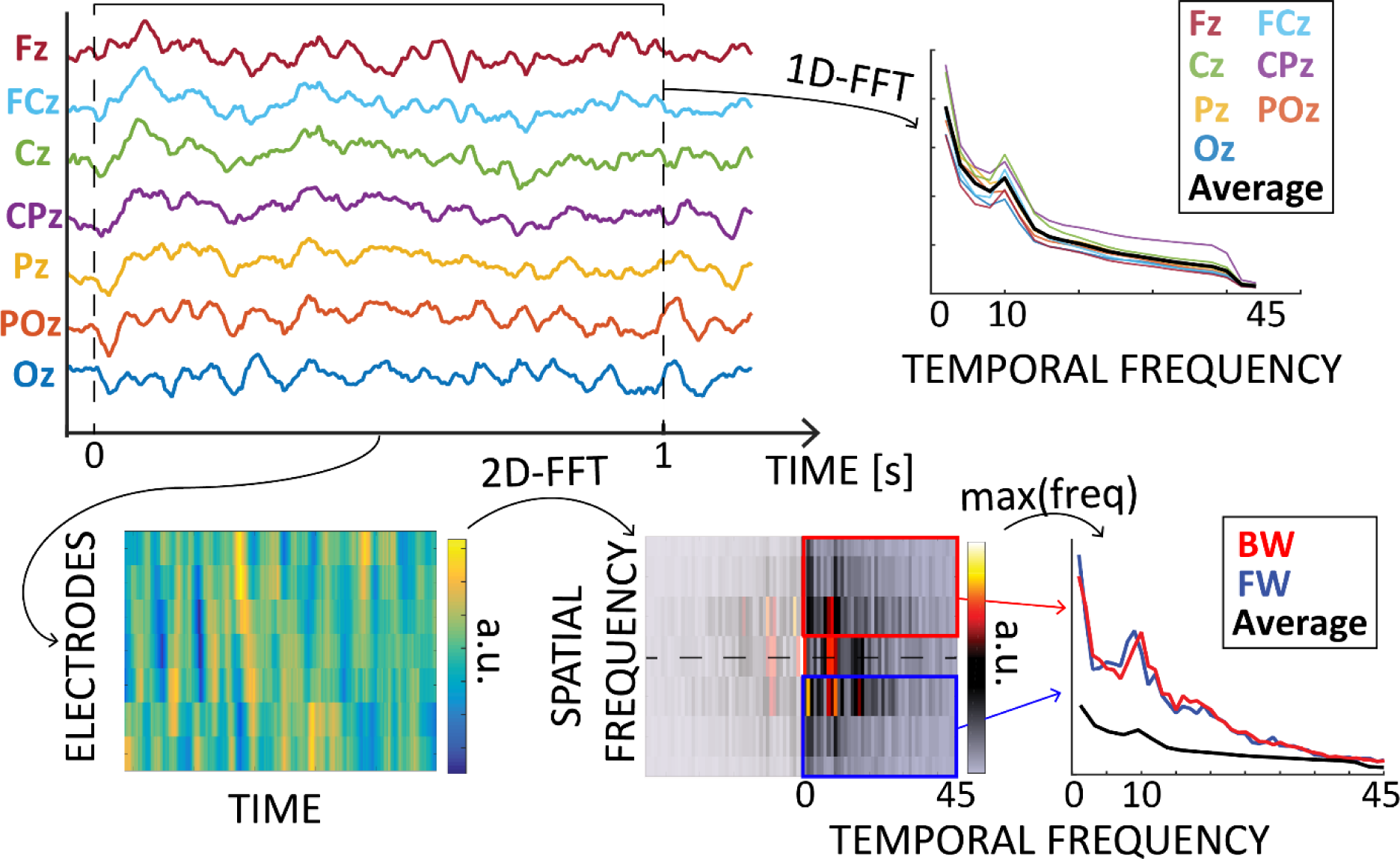
Quantifying traveling waves. From a 0.5 or 1-second time window, we extract 2D maps by combining seven midline electrodes. We then compute 2D-FFT to evaluate forward (FW – in blue) and backward (BW – in red) waves, as quantified by the maximum values in the lower- and upper-right quadrants, respectively. Lastly, we computed the waves amount in decibels by using the average 1D-FFT of each electrode as a baseline (see Methods).

For each of these images, we computed 2D Fast Fourier Transform (FFT) to quantify the amount of traveling waves propagating from occipital to frontal areas and vice versa (i.e., forward or backward, respectively). To determine an appropriate baseline (see Methods), we averaged each midline electrode 1D-FFT to obtain a baseline accounting for fluctuations in the overall power unrelated to the traveling waves (i.e., without the spatial information of the electrodes). The waves’ amount is expressed as a log-ratio in decibels [dB] (see Methods).

### Traveling waves during rest

In both the patients (N=121) and the control group (N=75), we quantified the spectral power in five frequency bands (θ: 3Hz - 7Hz, α: 8Hz – 12 Hz, low β:13Hz – 22Hz, high β: 23Hz –30 Hz, and γ: 30Hz – 45Hz) along the midline electrodes (from Oz to Fz, see figure 2). First, we compared each electrode’s spectral power between the two groups. For each frequency band, we performed Bayesian ANOVA, considering ELECTRODE and GROUP as factors (see Methods for details). As shown in figure 3A, we found a higher spectral power in the patients compared to the control group in the θ and α bands (GROUP factor BF_10_>>10^4^ in both frequency bands) but not in the beta- and gamma-bands. For all frequency bands, the ELECTRODE factor revealed a strong effect (all BF_10_>>10^66^). Next, we focused on the spatial component of brain oscillations by investigating differences in the spectra of traveling waves propagating forward (FW) and backward (BW). Figure 3B shows the spectra for both groups: as in our previous work (Alamia et al., 2023), in the control group, we found a typical spectral pattern with high backward waves in the α and low β (13Hz -22Hz) bands, and a flat profile in the forward waves (i.e., no difference between bands). We then compared the FW and BW spectra of the control group with those of the patients, considering the five frequency bands (θ, α, low and high β, and γ; figure 3B). We performed two-factor ANCOVAs, considering GROUP and BAND as factors and GENDER, AGE, and EDUCATION as covariates. For both FW and BW waves, we found a similar pattern of results: a very strong effect for the BAND factor (BF_10_>>10^11^ for both FW and BW waves) but mild to no effect in the GROUP factor (BW waves, BF_10_=7.519; FW waves, BF_10_=0.514); however, in both directions, we found a very robust effect of the GROUP x BAND interaction (for both FW and BW waves BF_10_>>400), revealing significant differences between the two groups. Concerning the covariates, there are no effects of EDUCATION or GENDER (BW waves, BF_10_<0.4; FW waves, BF_10_<0.8), but a small effect of AGE for the FW waves (BW waves, BF_10_=1.968; FW waves, BF_10_=4.684). We further analyzed these results by performing Bayesian ANOVAs for each frequency band separately, considering as factors GROUP and DIRECTION (either forward or backward).

**Figure 3.**
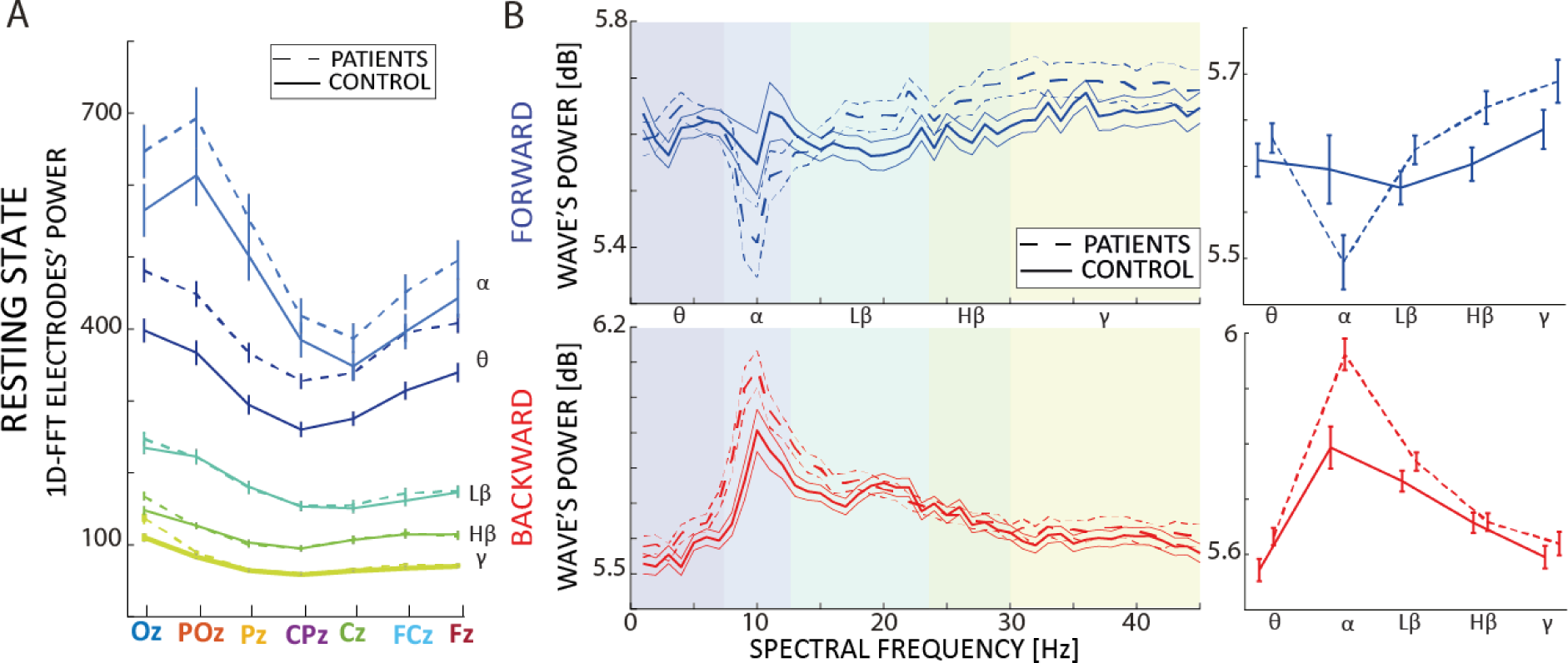
Differences in traveling waves between patients and controls during rest. A) Raw power for each spectral band in the midline electrodes (x-axis) for both the patients and control group in the resting state dataset. Each color represents a different frequency band. Error bars represent standard errors. B) The left panels illustrate the spectra for both forward (blue) and backward (red) waves for the two groups in both datasets; the right panels show the mean for each frequency band. Error bars represent standard errors.

Overall, the results show no difference between the groups (all frequency bands, 0.1<BF_10_<1), and a robust effect in the DIRECTION factor for θ, low β, and γ (all BF_10_>>10^7^). However, there was a strong interaction between GROUP and DIRECTION in the α band (BF_10_>>500), which, in line with our previous analysis, reveals distinct oscillatory dynamics between the patient and the control group. Specifically, these results confirm the difference shown in figure 3B: the most substantial effect was observed in the alpha-band (as confirmed by a larger Bayes Factor), where schizophrenia patients revealed an increase in backward waves and a decrease in forward waves.

### Traveling waves for backward masking

Previous results showed that the traveling wave pattern changes drastically between resting state EEG and visual evoked activity (Pang (庞兆阳) et al., 2020). In addition, according to the proposed Bayesian framework, schizophrenia patients have less precise priors at hierarchically lower sensory areas, thus weighing more the sensory information than healthy controls (Corlett et al., 2019; Sterzer et al., 2018). Accordingly, we expect an increase in alpha-band forward waves in patients following visual stimulation, in line with the hypothesis that alpha-band forward waves reflect precision-weighted sensory information (Alamia et al., 2020; K. J. Friston, 2019). For evoked activity, we analyzed a dataset of a Visual Backward Masking (VBM) task (see Herzog et al., 2004). In VBM, a briefly presented target is followed by a mask (figure 1C). There were 4 conditions: *Vernier only*, *Long SOA*, *Short SOA*, and *Mask only* conditions. In the *Vernier only* condition, there was no mask, whereas in the *Long SOA*, and *Short SOA* conditions, the target Vernier was followed by a mask with either an SOA of 150 or 30 ms, respectively. The *Mask only* condition provides a control condition, where no target was presented. As in the resting state analysis, we first assessed the spectral power in each electrode separately (figure 4A),. The Bayesian ANOVA, having ELECTRODE and GROUP as factors, revealed an overall higher power in the patient than in the control group in all frequency bands. As in resting state EEG, we also found a substantial effect of the ELECTRODE factor in all bands (all BF_10_>>10^48^). We then investigated the FW and BW traveling waves via two-factor ANCOVAs, with GROUP and BAND as primary factors and GENDER, AGE, and EDUCATION as covariates (figure 4B). We considered the waves after the onset of the visual stimulus before applying the baseline correction. Regarding the backward waves, we found a strong effect of BAND (BF_10_>10^18^), and moderate evidence for a difference between GROUP after the stimulus onset (BF_10_=8.47) but inconclusive before (BF_10_=0.911). We did not find evidence for an interaction (BF_10_<0.16) or the covariates (all 0.4<BF_10_<1.5). These results suggest that patients have a higher amount of backward waves irrespective of the frequency band after the onset of the target. Regarding the FW waves, we observed a strong effect in the BAND factor (BF_10_>10^8^) as well as in the GROUP factor (BF_10_>47). We found no evidence for the interaction (BF_10_<0.04), and inconclusive evidence for the covariate variables (all BF_10_<1.8). Next, we analyzed the changes in FW and BW waves with respect to the onset of the stimulus in each frequency band (i.e., applying a baseline correction computed on the 200ms before stimulus onset). In order to perform such analysis, we computed traveling waves in a time window ranging from 250ms before to 250ms after stimulus onset, with a temporal resolution of 100ms. Figure 4C illustrates the spectrogram for both forward and backward waves in both groups, whereas figure 4D gathers the same results for each frequency band separately.

**Figure 4.**
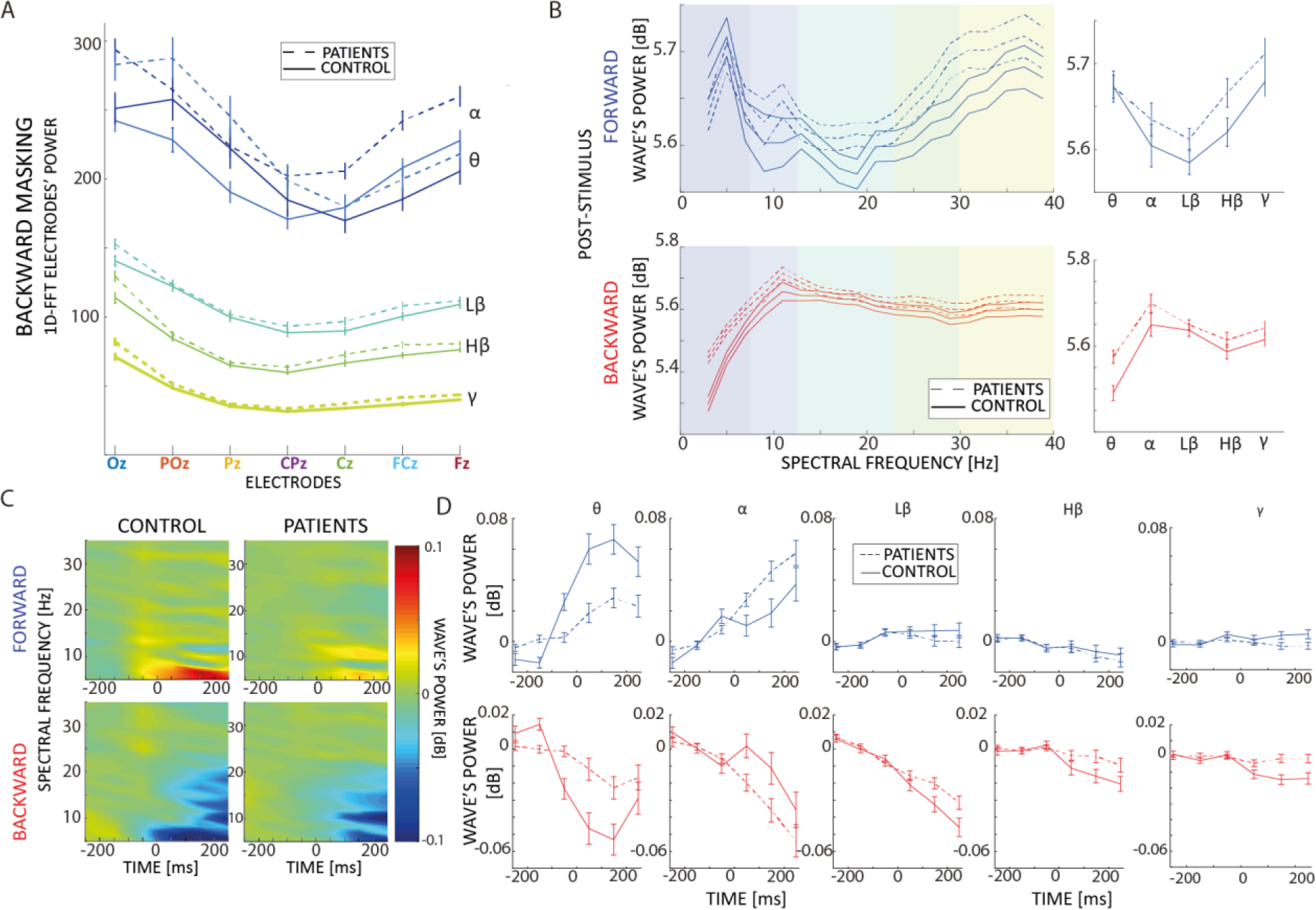
Differences in traveling waves between patients and controls during visual backward masking. A) Raw power for each spectral band in the midline electrodes (x-axis), for both the patients and control group in VBM dataset. Each color represents a different frequency band. B) The left panels illustrate the spectra for both forward (blue) and backward (red) waves for the two groups; the right panels show the mean for each frequency band. C) Spectrograms representing the forward and backward waves (upper and lower panels, respectively) for the control (left) and patient (right) groups. Color-coded values are baseline corrected considering 200ms before stimulus onset (at 0ms on the x-axis). D) Mean values and standard errors for each frequency band for the patient and control groups (dashed and solid lines, respectively); the x-axis represents time in milliseconds, with stimulus onset at 0ms. Error bars represent standard errors

Considering specifically alpha- and theta-band waves, our results reveal a difference between GROUP in the forward waves (*α*, BF_10_=21.52; *θ*, BF_10_=15.99), but with opposite effect: patients reveal an increase in alpha-band FW after stimulus onset, and vice versa in the theta-band. Regarding the BW waves, we found evidence for a difference in the theta-band (BF_10_>10^6^) but not in the alpha-band waves (BF_10_=0.13). Not surprisingly, in both FW and BW waves we found an effect of the TIME factor (*θ*: FW, BF_10_=145.48, BW, BF_10_>10^16^; *α*: FW, BF_10_=8.12, BW, BF_10_>10^14^), and an interaction between GROUP and TIME in all conditions (BF_10_>5 in all conditions) except in alpha BW waves (BF_10_=0.015). Considering beta and gamma-band traveling waves, as revealed in figure 4D, we did not observe any effect in the GROUP factor (all BF_10_<0.6) and an effect on TIME only for the BW low beta-band waves (BF_10_>10^16^), but not otherwise (all 0.1<BF_10_<1.8). All in all, these results confirm the prediction that the patient group show higher FW waves specifically in the alpha-band range, in line with the hypothesis that they rely more on precision-weighted sensory information.

Additionally, we investigated differences between groups in FW and BW alpha-band waves in each condition separately (i.e., *Vernier only*, *Long SOA*, *Short SOA*, and *Mask only* conditions, see above for details). As shown in supplementary figure S1, we confirm our results in all conditions except when only the mask was shown, corroborating the robustness of our results.

### Correlation between the resting-state and the VBM dataset

Out of the 144 patients and 96 control participants, 119 and 75, respectively, also belonged to the resting-state dataset, thus allowing us to reveal whether oscillatory traveling waves during rest are predictive of waves occurring during backward masking, reflecting general dynamics underlying predictive processes, which are not specific to the different tasks and/or experimental design.

We correlated band by band the amount of forward and backward waves in both groups (figure 5A), using Pearson correlations. In the visual backward masking dataset, we considered traveling waves before the onset of the stimulus to avoid the influence of visual processes (i.e., ERPs responses). As summarized in table 1, we found strong evidence for positive correlations between the two datasets for both forward and backward waves in all frequency bands and in both the control and the patient group.

**Table 1.** – Coefficients of correlation between resting states and backward visual masking traveling waves.

|  |  | PATIENTS |  | CONTROL |  |
| --- | --- | --- | --- | --- | --- |
| | | Pearson's r | $BF_{10}$ | Pearson's r | $BF_{10}$ |
| FORWARD | $\theta$ | 0.443 | 33457.916 | 0.376 | 30.532 |
| | $\alpha$ | 0.393 | 1775.222 | 0.504 | 4175.345 |
| | low $\beta$ | 0.510 | $3.781 \times 10^{+6}$ | 0.544 | 31240.309 |
| | high $\beta$ | 0.561 | $3.253 \times 10^{+8}$ | 0.589 | 456398.413 |
| | $\gamma$ | 0.607 | $3.341 \times 10^{+10}$ | 0.745 | $2.941 \times 10^{+11}$ |
| BACKWARD | $\theta$ | 0.403 | 2997.128 | 0.390 | 47.777 |
| | $\alpha$ | 0.391 | 1605.222 | 0.535 | 19551.076 |
| | low $\beta$ | 0.357 | 293.238 | 0.414 | 105.358 |
| | high $\beta$ | 0.406 | 3670.341 | 0.551 | 45799.009 |
| | $\gamma$ | 0.524 | $1.170 \times 10^{+7}$ | 0.663 | $9.435 \times 10^{+7}$ |

**Figure 5.**
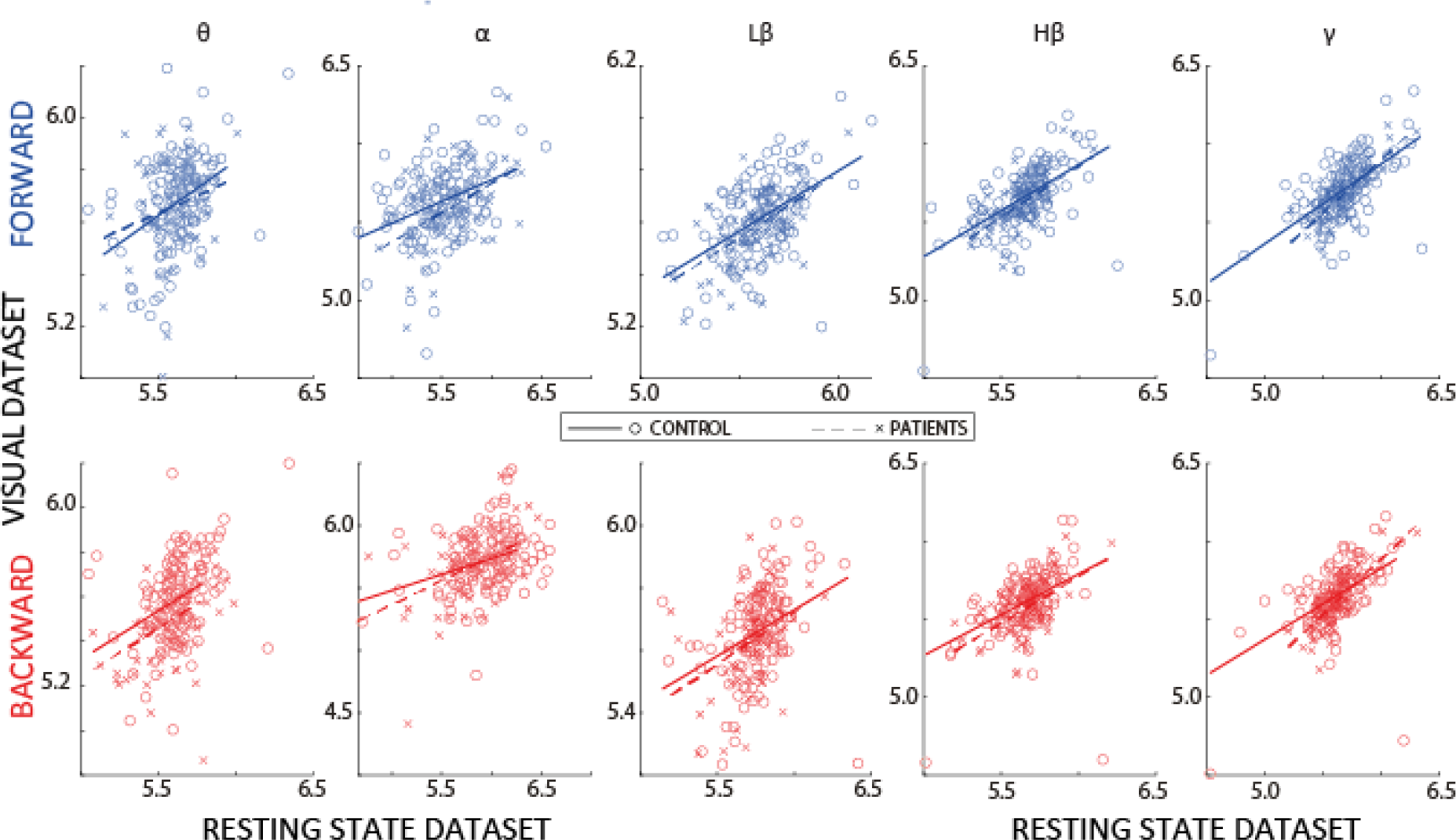
Correlations between datasets. A) Each subplot shows the correlation between traveling waves in the two datasets (resting state vs. VBM task). Columns represent different frequency bands, red and blue plots indicate backward and forward waves, respectively. The fitted line indicates significant (Pearson) correlations (see results for details).

### Testing the effects of medication and symptoms profile on traveling waves

To exclude potential confounds due to medication intake and to investigate whether traveling waves are associated to psychopathology in the schizophrenia patients, we correlated the amount of forward and backward waves in each frequency band with the Chlorpromazine (CPZ) equivalent dose that each patient received during the time of the recordings and with the scores in the negative and positive symptoms assessment (SANS and SAPS, respectively). All Pearson correlations in both datasets and for all frequency bands provided evidence in favor of the null hypothesis, that is, a lack of correlation between the amount of waves and CPZ-equivalents (resting state dataset: for both FW and BW waves in all frequency band -0.1<*r*<0.1, BF_10_<0.19 ; VBM dataset: for both FW and BW waves in all frequency band -0.14<*r*<0.1, BF_10_<0.22). Second, we performed a correlation between FW and BW waves and the SANS and SAPS scores (see Methods). We did not find evidence for correlations between traveling waves and SANS or SAPS scores (for both FW and BW waves in all frequency bands, - 0.13<*r*<0.13, BF_10_<0.23). We obtained similar results in the VBM dataset, in which we did not find any correlation in most frequency bands (-0.15<*r*<0.15, BF_10_<0.3), except for a small correlation between SAPS and BW waves in the low β (*r*=0.24, BF_10_=3.1), high β bands (*r*=0.26, BF_10_=5.16), and in the γ band (*r*=0.26, BF_10_=4.58); and between SAPS and FW waves in the γ band (*r*=0.27, BF_10_=7.404).

## Discussion

Analyzing EEG recordings in schizophrenia patients and healthy controls (da Cruz, Favrod, et al., 2020; da Cruz, Shaqiri, et al., 2020; Garobbio et al., 2021; Gordillo et al., 2023), we found direct evidence for a dysfunctional updating of beliefs about the world in schizophrenia patients (Fletcher & Frith, 2009). Previous studies have shown indirect evidence in favor of this interpretation (Corlett et al., 2009; Krystal et al., 2017). (Ellson, 1941; Kafadar et al., 2022; Kot & Serper, 2002). In our study, we targeted abnormalities in oscillatory traveling waves, which reflect the flow of information in predictive processes (Alamia et al., 2020; Alamia & VanRullen, 2019). Unlike previous work, our analysis allowed us to visualize and disentangle the different actors involved in the predictive coding process, flowing from higher to lower cortical areas, and prediction-errors, propagating in the opposite bottom-up direction in schizophrenia patients and healthy controls. Our results reveal a substantial increase in top-down and a decrease in bottom-up alpha-band traveling waves in schizophrenia patients compared to healthy participants in the resting state dataset, and the opposite pattern of results in the visual backward masking paradigm, demonstrating that schizophrenia patients have more precise priors (i.e., smaller variability) than healthy participants at hierarchically higher prior but less precise priors in lower sensory areas (Corlett et al., 2019; Friston, 2005; Powers et al., 2017). Importantly, these findings describe not only the temporal but also the spatial component of brain rhythms, supporting the key idea that oscillatory dynamics are best understood when interpreted as traveling waves propagating across cortical regions, coordinating and synchronizing different brain regions (Alexander et al., 2009; Muller et al., 2018).

Our results may reconcile previously contradictory findings. Some authors proposed that positive symptoms in schizophrenia relate to abnormalities in prior expectations (Wacongne, 2016; Weilnhammer et al., 2020); on the other hand, other authors argued against this interpretation, based on indirect experimental findings, such as intact illusion perception and intact contextual processing (Choung et al., 2022; Grzeczkowski et al., 2018; Lhotka et al., 2023). To reconcile these contradicting findings, a more nuanced framework was proposed to account, separating priors in the lower areas from those in higher ones (Corlett et al., 2019; Sterzer et al., 2018). For example, visual illusions may rely on relatively lower-level priors, which affect visual perception specifically, whereas schizophrenia patients may have impairments in higher-level priors, involved in higher-order functions. Accordingly, schizophrenia patients proved to be more sensitive than controls to illusions involving higher-order processing, such as temporal expectations in the triple flash illusion (Norton et al., 2008), than simpler visual illusions. Our results are in line with this hypothesis. In particular, we measured traveling waves during rest and during a visual backward masking task, in which neither predictions nor sensory expectations play any substantial role (especially in resting state), suggesting that alpha-band traveling waves do not reflect specific perceptual features of the task, but rather broader brain states. This conclusion is further corroborated by the lack of difference between distinct conditions in the visual backward masking task (fig. S1).

Neural oscillations have long been studied in schizophrenia research and have been proposed to play a crucial role in coordinating neural activity, and aberrancies in their strength and synchronization may be a core pathophysiological mechanism of schizophrenia (Uhlhaas & Singer, 2006, 2010). Our findings are consistent with previous studies, in which differences between schizophrenia patients and healthy controls have been observed in several frequency bands during resting-state (Gordillo et al., 2023; Newson & Thiagarajan, 2019) and different tasks or experimental conditions (Hirano & Uhlhaas, 2021; Javitt et al., 2018; Roach & Mathalon, 2008). For example, during resting-state, schizophrenia patients show increased activity in the delta, theta and beta bands (Venables et al., 2009), whereas activity in the alpha and gamma bands is strongly decreased, compared to healthy controls (Knyazeva et al., 2008; Uhlhaas & Singer, 2013). Similarly, during visual processing, activity in the delta, theta and gamma bands has been shown to be reduced in schizophrenia patients (Martínez et al., 2018; Uhlhaas et al., 2006). Accordingly, our results reveal consistent differences between schizophrenia patients and healthy participants in a broad-band fashion but with a more substantial effect in the alpha-band, consistent with our hypothesis. As in our work, previous studies also related brain oscillations to predictive coding, associating prediction and prediction-error to different frequency bands based on consistent experimental evidence (Bastos et al., 2012, 2015; Michalareas et al., 2016; Vezoli et al., 2021). Specifically, gamma oscillations (>30Hz) reflect local cortical processes, and besides characterizing a wide range of cognitive functions (Lundqvist et al., 2016; Zhang et al., 2012), they proved to match reliably sensory expectations and prediction errors, corroborated by the fact that the regular repetition and the unexpected omission of a stimulus respectively decrease and increase gamma oscillations’ activity (Fujioka et al., 2009; Iversen et al., 2009). In agreement with our interpretation, alpha and beta band oscillations (∼8-30Hz) have been related to top-down activity, carrying predictions from higher to lower brain regions (Haegens et al., 2011; Samaha et al., 2015; van Pelt et al., 2016). In our results, the relationship between top-down traveling waves and prior belief is further corroborated by the moderate positive correlation between backward waves in the beta and gamma bands, and positive symptoms assessment (SAPS), which quantifies the symptoms related to hallucinations, delusions, and aberrancies in perception. Importantly, our study is the first one to consider oscillations as traveling waves propagating through the cortex in a forward and backward manner, thus, taking into account not only the temporal but also the spatial dimension of oscillations. It is this aspect that allowed us to reconcile the mixed findings about Bayesian predictions in schizophrenia.

All in all, our findings demonstrate that schizophrenia patients have stronger high-level priors, which elicit stronger alpha-band oscillations, and weaker low-level priors, leading to an overall dysfunctional updating of their cognitive and sensory world model. Our results provide direct and convincing evidence in favor of hierarchical-specific abnormalities in the prior of schizophrenia patients.

## Methods

### Participants

We present resting-state and evoked EEG data collected from two groups of participants: schizophrenia patients (N=146) and healthy controls (N=96). Resting-state EEG data were recorded for 121 schizophrenia patients and 75 healthy controls. Visual backward masking (VBM) EEG data were recorded from 144 schizophrenia patients and 96 healthy controls, with 119 schizophrenia patients and 75 healthy controls also having resting-state recordings. Previous studies have already utilized the resting-state data for 121 schizophrenia patients and 75 healthy controls (Gordillo et al., 2023), as well as the VBM EEG data for 121 schizophrenia patients and 94 healthy controls (Garobbio et al., 2021).

Schizophrenia patients were recruited from the Tbilisi Mental Health Hospital or the psycho-social rehabilitation center. Among the patients, 49 were inpatients, while 97 were outpatients. Diagnostic assessment for patients was determined using the Diagnostic and Statistical Manual of Mental Disorders Fourth Edition (DSM-IV) through interviews, information from staff, and examination of patients’ records. The severity of the patient’s symptoms was assessed by an experienced psychiatrist using the Scale for the Assessment of Negative Symptoms (SANS) and the Scale for the Assessment of Positive Symptoms (SAPS). Out of the 144 patients, 131 were receiving neuroleptic medication. The equivalent doses of Chlorpromazine (CPZ) are provided in Table 1.

Controls were selected from the general population in Tbilisi, aiming to closely match the demographics of the patient group. All control participants had no psychiatric axis I disorders and no family history of psychosis. Exclusion criteria included alcohol or drug abuse, severe neurological incidents or diagnoses, developmental disorders such as autism spectrum disorder or intellectual disability, or other significant somatic illnesses affecting mental functioning. These criteria were assessed through interviews conducted by certified psychiatrists. Detailed characteristics of the groups are presented in Table 1.

Before participating in the study, all individuals provided informed consent and were informed of their right to withdraw from the study at any time. The study procedures were conducted in accordance with the Declaration of Helsinki (except for preregistration) and were approved by the Ethical Committee of the Institute of Postgraduate Medical Education and Continuous Professional Development in Georgia (protocol number: 09/07; title: "Genetic polymorphisms and early information processing in schizophrenia").

**Table 1.**

|  | VBM patients | VBM controls | Statistics | RS patients | RS controls | Statistics |
| --- | --- | --- | --- | --- | --- | --- |
| Gender (M/F) | 115/29 | 49/47 | $\chi^2(1)=22.108$, $P<0.001^a$ | 22/99 | 39/36 | $\chi^2(1)=24.702$, $P<0.001^a$ |
| Age (years) | 35.5±9.1 | 35.0±8.3 | $t(238)=-0.393$, $P=0.695^b$ | 35.8±9.2 | 35.1±7.7 | $t(194)=-0.519$, $P=0.604^b$ |
| Education (years) | 13.2±2.6 | 15.2±2.9 | $t(238)=-5.432$, $P<0.001^b$ | 13.3±2.6 | 15.1±2.9 | $t(194)=4.418$, $P<0.001^b$ |
| Handedness (L/R) | 7/137 | 6/90 | $\chi^2(1)=0.217$, $P=0.641^a$ | 6/115 | 4/71 | $\chi^2(1)=0.013$, $P=0.908^a$ |
| Illness duration (years) | 10.7±8.5 |  |  | 10.8±8.7 |  |  |
| SANS | 10.1±5.2 |  |  | 10.1±5.2 |  |  |
| SAPS | 8.7±3.2 |  |  | 8.6±3.2 |  |  |
| CPZ equivalent | 578.3±391.6 |  |  | 561.1±389.4 |  |  |
| Visual acuity | 1.4±0.4 | 1.6±0.4 | t(238)=-3.388, P<0.001<sup>b</sup> | 1.4±0.4 | 1.6±0.4 | t(194)=2.969, P=0.003<sup>b</sup> |
| Vernier duration* | 20 [20, 50] | 20[20, 20] | X<sup>2</sup>(1)=37.949, P<0.001<sup>c</sup> | 20 [20, 52.5] | 20[20, 20] | X<sup>2</sup>(1)=28.25, P<0.001<sup>c</sup> |
| Note: SANS, Scale for the Assessment of Negative Symptoms; SAPS, Scale for the Assessment of Positive Symptoms; CPZ, Chlorpromazine equivalents<br><sup>a</sup> Pearson's chi-squared test<br><sup>b</sup> Two-sided independent samples t-test<br><sup>c</sup> Mood's median test<br>*Median [25<sup>th</sup> percentile, 75<sup>th</sup> percentile]; Vernier duration value is missing for one patient in both datasets |  |  |  |  |  |  |

### Experimental procedure

#### Visual backward masking

##### Stimuli

The stimuli consisted of two vertical line segments of 10’ (arc-minutes) length with a gap of 1’. The lower bar was slightly offset either to the left or right compared to the upper bar. The offset was fixed at about 1.2’. The mask was composed of 25 verniers without offset separated by 3.33’.

##### Apparatus

The stimuli were displayed on a cathode ray tube screen (Siemens Fujitsu P796-1) with a refresh rate of 100 Hz. The screen resolution was 1024 x 768 pixels. Participants were seated at 3.5m from the monitor in a dimly lit room. A pixel comprised about 18’’ (arc-seconds). Stimuli were white. The luminance was 100 cd/m^2^ (measured with a Gretag Macbeth Eye-One Display 2 colorimeter) on a black background of < 1 cd/m^2^.

##### Adaptive procedure

Further details of the paradigm can be found in a previous study (Herzog et al., 2004). We determined the Vernier Discrimination Threshold (VD) required for participants to achieve 75% of correct responses in identifying a vernier offset of 0.6’. Participants were required to achieve a VD shorter than 100 milliseconds. The vernier stimulus, with the individualized VD for each participant and an offset of 1.2’, was presented, followed by an interstimulus interval and a mask lasting 300 milliseconds. Using a staircase procedure, we adaptively determined the target-mask stimulus onset asynchrony (SOA), calculated as the sum of VD and interstimulus interval (ISI), to achieve a performance level of 75% correct responses. This was done using Parametric Estimation by Sequential Testing (PEST). Each participant completed the test twice, and the results of the first and second tests were averaged and are presented in Table 1.

##### EEG experiment

ERP latencies and amplitudes vary with VD. Hence, for the EEG experiment, we maintained VDs and SOAs constant thereby using the same stimuli for all observers. We set the VD to 30 milliseconds, which corresponds to the average VD observed in previous studies involving patients (Chkonia et al., 2010; Herzog et al., 2004). Our experiment encompassed four distinct stimulus conditions (Favrod et al., 2017, 2019; Plomp et al., 2013). In the *Vernier Only* condition, only the target vernier stimulus was presented. The *Long SOA* condition involved the presentation of the mask following the target vernier, with an SOA of 150 milliseconds. In contrast, the *Short SOA* condition featured the immediate presentation of the mask after the target vernier, resulting in an SOA of 30 milliseconds. The selection of SOAs for the *Long SOA* and *Short SOA* conditions was based on the mean SOAs observed in previous studies involving both schizophrenia patients and controls (Chkonia et al., 2010; Favrod et al., 2018; Herzog et al., 2004; Plomp et al., 2013). To provide a control condition, we included the Mask Only condition, wherein only the mask stimulus was presented. In this particular case, accuracy was determined by comparing the left/right offset response to a randomly chosen hypothetical offset.

##### Resting-state

Resting-state EEG data were recorded for 5min. Participants were seated in a dimly lit room and were instructed to close their eyes and relax during the recording.

### EEG recording and Preprocessing

EEG was recorded using a BioSemi Active 2 system with 64 Ag-AgCl sintered active electrodes, referenced to the common mode sense electrode. The sampling rate was 2048 Hz. Offline data were downsampled to 512 Hz and preprocessed using an automatic preprocessing pipeline (da Cruz et al., 2018). For resting-state EEG data, the preprocessing included the following steps: high-pass filtering at 1 Hz; power line noise removal (using CleanLine; www.nitrc.org/projects/cleanline); re-referencing to the bi-weight estimate of the average of all electrodes; removal and 3D spline interpolation of bad electrodes; removal of bad epochs; independent component analysis (ICA) to remove artifacts related to eye-movements, muscle activity, and bad electrodes, and removal of epochs with artifacts (1s epochs). For VBM EEG data, the preprocessing was similar to the resting-state data with the difference that a band-pass filtering between 1 and 40 Hz was performed instead of high-pass filtering at 1 Hz. VBM EEG data from 2 patients (both present in the resting-state dataset) and 2 healthy controls (1 present in the resting-state dataset) were discarded due to excessive EEG artifacts.

### Traveling wave analysis

As in our previous studies (Alamia et al., 2020; Pang (庞兆阳) et al., 2020) we quantified traveling waves’ propagation along seven midline electrodes, running from occipital to frontal regions (Oz, POz, Pz, CPz, Cz, FCz, Fz, as shown in figure 1). After segmenting the signal in a 1-second time window (and 500ms for the VBM dataset), we stacked the signals from the seven electrodes to create 2D maps, with time and electrodes as axes. From each map, we compute 2D-FFT transformation: importantly, the power in the lower and upper quadrants quantifies the amount of forward (FW - from occipital to frontal electrodes) and backward (BW - from frontal to occipital) waves, respectively (see figure 1). For each frequency in the 2-45Hz range, we considered the maximum values in both the upper and lower quadrant, obtaining a spectrum for both *BW* and *FW* waves, respectively. We then needed to determine a baseline to account for fluctuations in the overall power unrelated to the traveling waves. For this reason, we computed the *average* 1D-FFT of each midline electrode, which provides a baseline accounting for the spectral power unrelated to the traveling waves (i.e., without the spatial information obtained by combining the electrodes). We finally obtained the waves’ amount in decibels [dB] as:

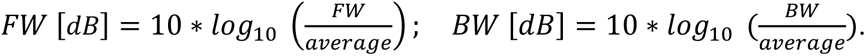

This value quantifies the total waves compared to the null distribution, thus being informative when contrasted against zero. It is important, however, to notice that this is a surface-level analysis, and it is not informative about the underlying sources. When interpreting these traveling waves results, it is also important to consider issues related to long-range connections and distortions due to scalp interference (Alexander et al., 2019; Nunez, 1974). In particular, when considering traveling waves analysis with non-invasive surface recordings (such as EEG), it is crucial to consider distortions and interference between coincident waves and limitations due to the electrode configurations (Alexander et al., 2019; Nunez, 1974, 2000). Specifically, it’s possible to detect only waves shorter than the spatial length of the sensor array and waves longer than twice the distance between electrodes (due to the Nyquist criterion in space). In addition, different cortical processes may generate a similar pattern of traveling waves visible via surface recordings (Alamia & VanRullen, 2023), thus limiting the interpretation of possible underlying sources.

### Statistical analyses

We analyzed frequency-band averaged traveling waves by means of Bayesian ANOVAs. In all analyses, we computed Bayes Factors (BF) as the ratio between the models testing the alternative against the null hypothesis. All BFs are denoted as BF_10_ throughout the paper. In practice, BFs provide substantial (BF>∼5) or strong (BF>∼10) evidence in favor of the alternative hypothesis, and low BF (BF<∼0.5) suggests a lack of effect (Masson, 2011). In each dataset, we performed a Bayesian ANOVA on each electrode spectral power considering ELECTRODE (from Oz to Fz along the midline) and GROUP (patients and control) as fixed factors and SUBJECT as the random term. We also performed an ANCOVA considering GENDER, AGE, and EDUCATION as covariates, GROUP and BAND as fixed terms, and SUBJECT as random terms. Lastly, all Bayesian correlations were computed considering both Pearson and Kendall, but we reported for simplicity only Bayes Factor for the Pearson *r*, (similar results were obtained considering Kendall correlations). We performed all analyses in JASP (Love et al., 2019), considering default uniform prior distributions.

## Acknowledgment

This project was funded by the European Union under the European Union’s Horizon 2020 research and innovation program (grant agreements No. 101075930 to Andrea Alamia) and the National Centre of Competence in Research (NCCR) Synapsy financed by the Swiss National Science Foundation under grant 51NF40-185897. Views and opinions expressed are however those of the author(s) only and do not necessarily reflect those of the European Union or the European Research Council (ERC). Neither the European Union nor the granting authority can be held responsible for them

---

**Figure S1.**
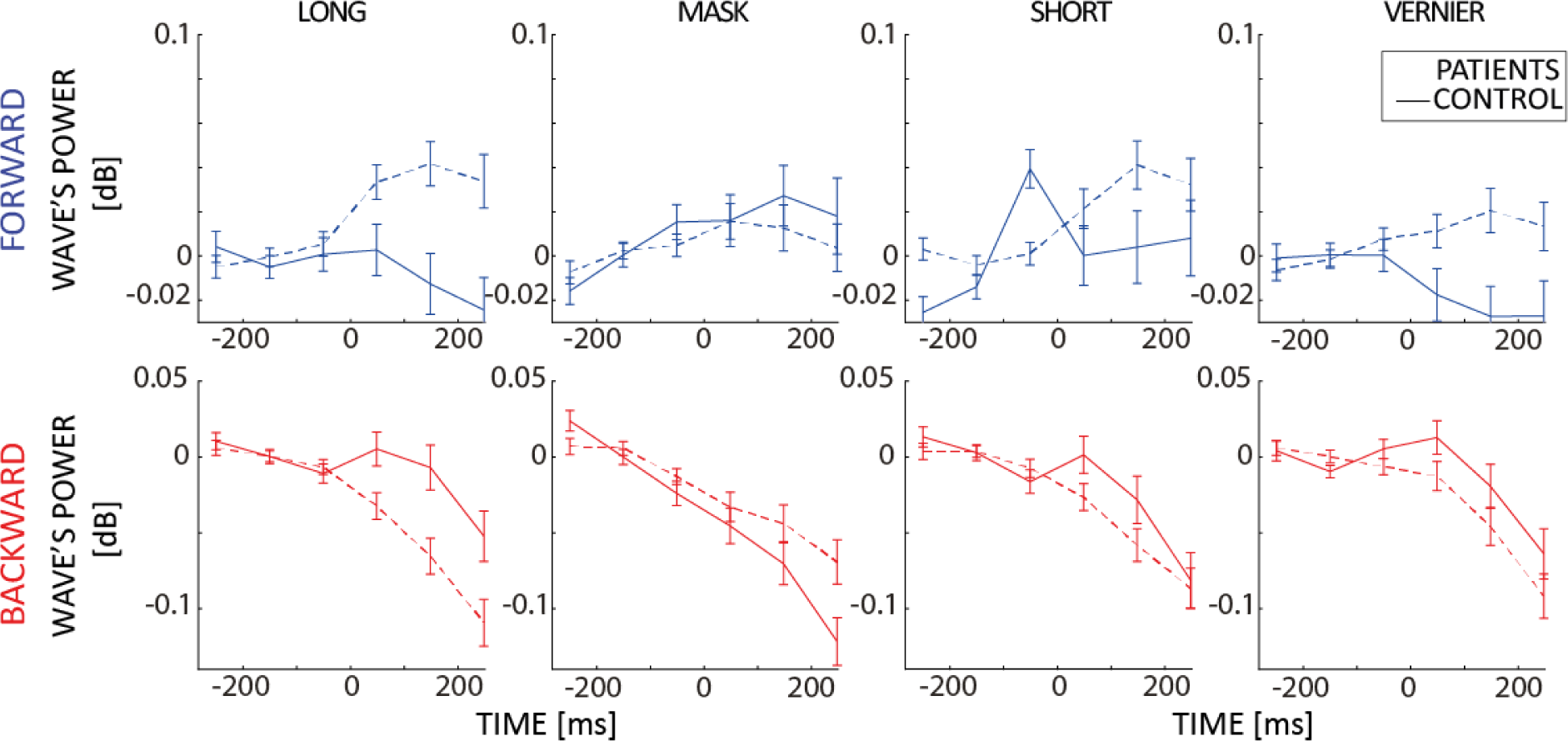
Differences in alpha-band traveling waves between patients and control in the visual dataset task. Each row represents a different condition. As in the main figures, each subplot shows the mean values and standard errors for the patient (dashed lines) and the control groups (solid lines) before and after the onset of the stimulus at 0ms.

